# Oncogenic c-Myc activity increases tolerance to proteotoxic and genotoxic stress through regulation of HSF1

**DOI:** 10.1101/2022.02.02.478796

**Authors:** Joe Jones, Koshiro Kiso, Cosetta Bertoli, Robertus A. M. de Bruin

## Abstract

Oncogenes, such as c-Myc, enhance growth and proliferative signaling to promote continuous cell cycle divisions, the hallmark of cancer. The inadvertent consequence of this is an increase in cellular stresses. However, whether and how oncogenes can directly contribute to cellular stress tolerance, and how much cancer cells rely on these mechanisms for survival, remains poorly understood. Here we show that c-Myc-dependent proteotoxic stress contributes to the generation of genotoxic stress. We reveal an important role for the transcription factor Heat Shock Factor 1 (HSF1) in the tolerance to both these c-Myc-induced stresses. c-Myc upregulates HSF1 directly, by activating its expression, and indirectly, via c-Myc-dependent proteotoxic stress activation. In addition to relieving c-Myc-induced proteotoxic and genotoxic stress, HSF1 also enables DNA damage response signalling through *γ*H2AX. Consequently, acute depletion of HSF1 significantly increases c-Myc-driven genome instability and decreases cell viability. Our results establish that c-Myc-dependent regulation of HSF1 ensures that proteotoxic and genotoxic stress, resulting from c-Myc-induced enhanced growth and proliferation, are compatible with cell survival.

## Introduction

Cancer pathogenesis involves a series of mutagenic events that drive enhanced growth and continued proliferation. Whilst this allows for the classical hallmarks of cancer, such as an unlimited proliferation potential and the ability to evade apoptotic signals, it also leads to the stress phenotypes of cancer (1–3), such as proteotoxic and genotoxic stress. The consequential rise in oncogenesis-related stresses increases the dependency of cancer cells on stress support genes and/or pathways for their survival, also known as non-oncogene addiction.

Proteotoxic stress, an event that disturbs proteostasis, is one of the most common cancer stress phenotypes (3). Cancer cells experience proteotoxic stress as a consequence of increased protein translation rates, required to satisfy the anabolic demands of transformation and proliferation (4, 5); harsh tumour microenvironments that perturb proteostasis (3, 6); and an increase in aneuploidy that promotes protein aggregation (7, 8). Tolerance to proteotoxic stress largely depends on the transcription factor Heat Shock Factor 1 (HSF1), which mediates a transcriptional response by pro-moting transcription of heat shock proteins (HSPs) that guide protein folding, trafficking and disaggregation, thereby lowering proteotoxic stress levels (9, 10). Consequently, cancers have an increased reliance on the HSF1-mediated proteotoxic stress response, with knockout of *Hsf1* in mouse models of cancer impairing tumourigenesis (11, 12). Oncogenic signalling is thought to be able to upregulate HSF1 directly (13–19) to facilitate an increased tolerance to proteotoxic stress and other cancer-related stresses, but our understanding of this is incomplete.

HSF1 has been linked to a variety of biological processes distinct from the proteotoxic stress response (20–22). Of these the most notable is the DNA damage response (DDR) (23, 24), which is required for the tolerance to genotoxic stress, another cancer stress phenotype. Whilst the exact role of HSF1 in the DDR, and the molecular mechanisms involved, remain largely unknown, it places HSF1 at the nexus of the tolerance to two of the main cancer stress phenotypes. Oncogenic c-Myc activity, a frequent event in multiple human malignancies (25), causes intrinsic proteotoxic stress due to enhanced protein synthesis to drive rapid cell growth and proliferation (26). Sustained proliferative signalling is thought to be at the basis of c-Myc-dependent genotoxic stress (27), driving genomic instability. However, whether genomic instability is also driven in part through an increase in proteotoxic stress remains underexplored. Overall, the mechanisms enabling adaptation to these c-Myc-induced stress are not fully understood.

Here, we report how oncogenic c-Myc supports tolerance to increased proteotoxic and genotoxic stress through regulation of HSF1. We find that c-Myc-induced proteotoxic stress contributes to the generation of DNA damage. c-Myc directly upregulates HSF1 via transcriptional induction and indirectly through c-Myc-induced proteotoxic stress. We show that, in addition to its central role in proteotoxic stress tolerance, HSF1 is required for a functioning DDR, through the stabilisation of newly phosphorylated *γ*H2AX protein, required for the first step in recruiting and localizing DNA repair proteins (28, 29). Consequently, c-Myc increases adaptation to proteotoxic and genotoxic stress through activation of HSF1.

## Results

### c-Myc-induced proteotoxic stress generates DNA damage

We used immortalised retinal pigment epithelial (RPE1) cells stably expressing c-Myc conjugated to the oestrogen receptor (ER), hereafter c-Myc-ER cells, to investigate the consequences of c-Myc activation. Activation of c-Myc is induced upon addition of 4-hydroxytamoxifen (4-OHT) to cell culture media and RPE1 cells expressing an unconjugated ER protein, hereafter ER-Empty cells, are used as a control (Supplementary Figure S1a,b). Our c-Myc inducible system has been extensively characterized and shown to cause proliferation, DNA damage and genomic instability upon induction (30). To test if acute activation of c-Myc leads to proteotoxic stress, protein aggregation levels were assessed in c-Myc-ER cells following activation of c-Myc for the indicated timepoints, before fixing cells and staining with the fluorescent dye Proteostat, which stains protein aggresomes (31). c-Myc activation leads to acute accumulation of protein aggregates over time, peaking at 24h (Figure 1a). Protein aggregation following 24h of c-Myc activation is at a similar level as the positive control, where cells were treated with the proteasome inhibitor bortezomib (BTZ) for 6h. Addition of 4-OHT to ER-Empty cells does not cause an increase in proteostat staining (Supplementary Figure S1c). These data show that in our system c-Myc activation generates proteotoxic stress.

**Fig. 1.**
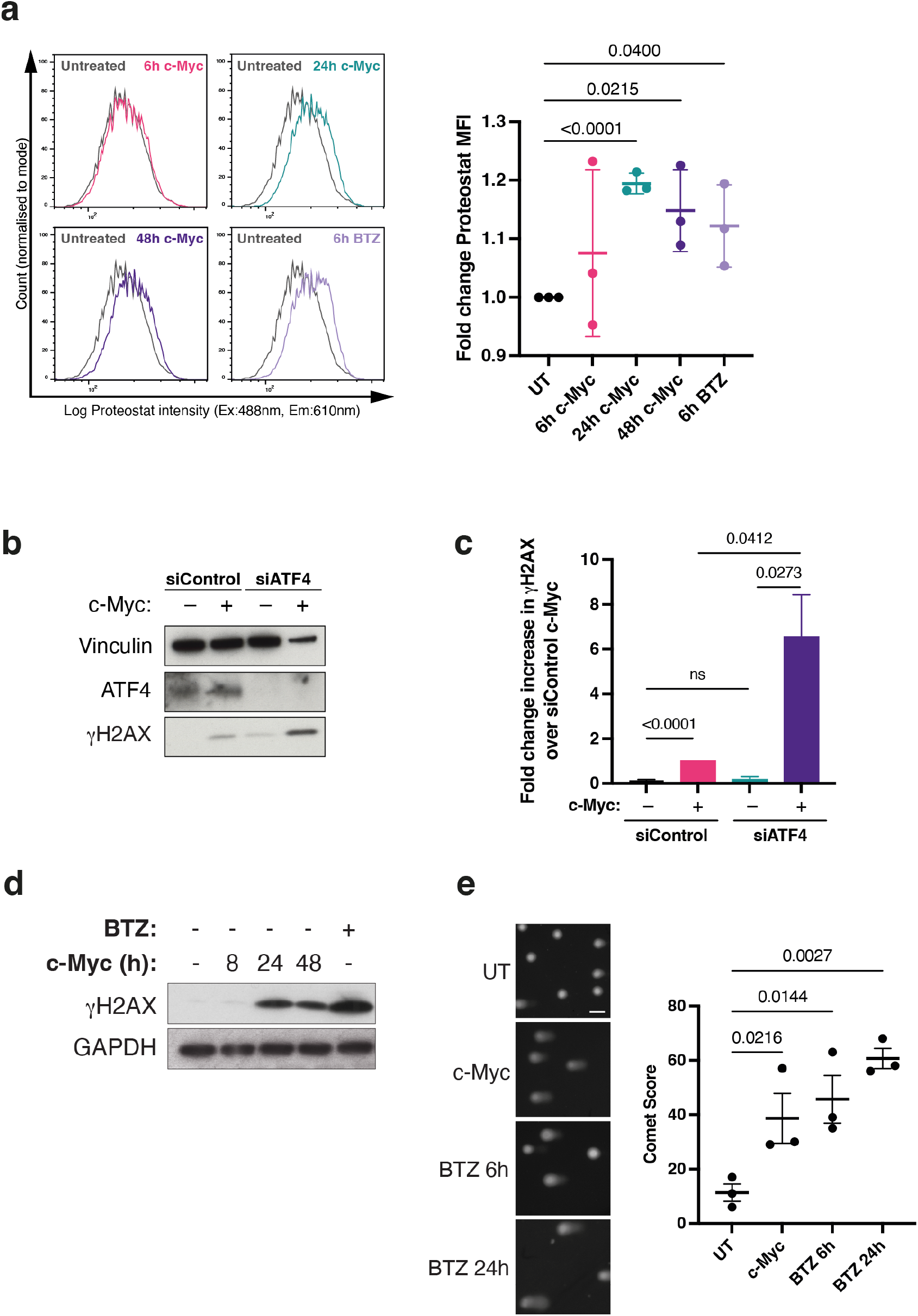
c-Myc-induced proteotoxic stress generates DNA damage. **(a)** c-Myc-ER cells had c-Myc activated for the indicated timepoints, or bortezomib (BTZ) treatment for 6h, before cells were fixed and stained with Proteostat for flow cytometry analysis, representative histograms shown. Fold change proteostat median fluorescent intensity (MFI) was calculated for each sample, n=3, error bars are SEM. Unpaired Student’s t-test was used for statistical analysis. **(b)** c-Myc-ER cells transfected with siControl or siATF4 siRNAs were untreated or had c-Myc activated for 24h, before protein lysates were collected for Western blot. Representative blot shown from n=3 independent experiments. **(c)** Fold change increase in *γ*H2AX protein levels over siControl c-Myc cells for each sample, error bars are SEM, n=3. Unpaired Student’s t-test used for statistical analysis. **(d)** c-Myc-ER cells had c-Myc activated for the indicated timepoints or BTZ treatment for 6h before protein lysates were collected for Western blot. Representative blot shown from n=3 independent experiments. **(e)** c-Myc-ER cells had c-Myc activated for 24h or BTZ treatment for the indicated timepoints before cells were collected for alkaline comet assay analysis. Scale bar = 50mm. Comet score for each condition shown, n=3, error bars are SEM. Unpaired Student’s t-test was used for statistical analysis.

Oncogenic c-Myc activation is known to generate DNA damage, thought mostly through the generation of replication stress-induced DNA damage (27). We wondered if proteotoxic stress also contributes to c-Myc-induced DNA damage generation. To test this, we sought to increase c-Myc-induced proteotoxic stress levels by increasing protein translation through knockdown of ATF4, to establish if this increases DNA damage levels. ATF4 is required to increase 4E-BP1 activity, which is negatively regulated by mammalian target of rapamycin complex 1 (mTORC1)-dependent phosphorylation, to relieve MYC-induced proteotoxic stress (32– 34). We confirm that ATF4 knockdown causes an increase in protein aggregation levels in our c-Myc inducible system, as assessed by Proteostat staining (Supplementary Figure S1d). Next, we established if increased c-Myc-induced proteotoxic stress, by ATF4 knockdown, affects c-Myc-induced DNA damage levels, via quantification of hyperphosphorylated H2AX (*γ*H2AX) protein levels. *γ*H2AX is one of the earliest markers of ATM-mediated DNA damage checkpoint activation and is therefore a quantitative and sensitive marker of DNA damage levels (28, 29). c-Myc activation causes an increase in *γ*H2AX protein levels, indicative of c-Myc-induced DNA damage (Supplementary Figure S1e), which is markedly increased upon ATF4 knockdown (Figure 1b,c). Together, these data suggest that c-Myc-induced proteotoxic stress contributes to the accumulation of c-Myc-induced DNA damage.

To establish if proteotoxic stress per se can cause the accumulation of DNA damage, we induced proteotoxic stress via treatment with the proteasome inhibitor bortezomib (BTZ) for 6h and assessed *γ*H2AX levels alongside c-Myc-induced *γ*H2AX levels (Figure 1d). BTZ has previously been described to increase DNA damage following irradiation treatment in HeLa cells (35) and in sensory ganglia neurons (36).

Our data show that BTZ-induced proteotoxic stress alone causes DNA damage to a comparable level to, or even greater extent than, c-Myc-induced DNA damage levels. To rule out that increased *γ*H2AX levels are a consequence of protein accumulation effects caused by inhibition of the proteasome, we assayed DNA damage directly using the comet assay. First, we show that the comet assay can reliably detect c-Myc-induced DNA damage by treatment of c-Myc-ER and ER-empty cells with or without 4-OHT (Supplementary Figure S1f). Next, we found that 6h and 24h of BTZ treatment led to a dose-dependent increase in DNA damage, to similar or even greater extent than c-Myc activation (Figure 1e). Together, this suggests that proteotoxic stress can generate DNA damage and that c-Myc-induced DNA damage may be driven in part by proteotoxic stress. This puts the response to proteotoxic stress at the centre of both the tolerance to proteotoxic stress itself but also the prevention of genotoxic stress, two main cancer stress phenotypes.

### HSF1 is upregulated upon c-Myc activation post-translationally and transcriptionally

One of the major proteotoxic stress response pathways within the cell is the heat shock response, which is mediated by the transcription factor HSF1. Upon a rise in proteotoxic stress, HSF1 is activated through post-translational modification and deploys its extensive transcriptional network to restore proteostasis. As we have shown that c-Myc activation causes a rise in proteotoxic stress, we next ask if the HSF1-dependent response is induced in response to c-Myc activation. Levels of HSF1 phosphorylation at serine 326, required for HSF1’s transcriptional activity (37), increase upon activation of c-Myc, peaking at 24h of 4-OHT treatment (Fig. 2a). In addition, HSF1 must translocate from the cytoplasm into the nucleus in order to induce its transcriptional network (38). We show that c-Myc activation causes an increase in the nuclear localisation of pHSF1 S326, with 24h of c-Myc activation causing nuclear localisation to the same extent as 6h BTZ treatment (Fig. 2b). Together, these data show that c-Myc promotes the post-translational modification and nuclear accumulation of HSF1, required for its activation, likely as the result of c-Myc-induced proteotoxic stress.

**Fig. 2.**
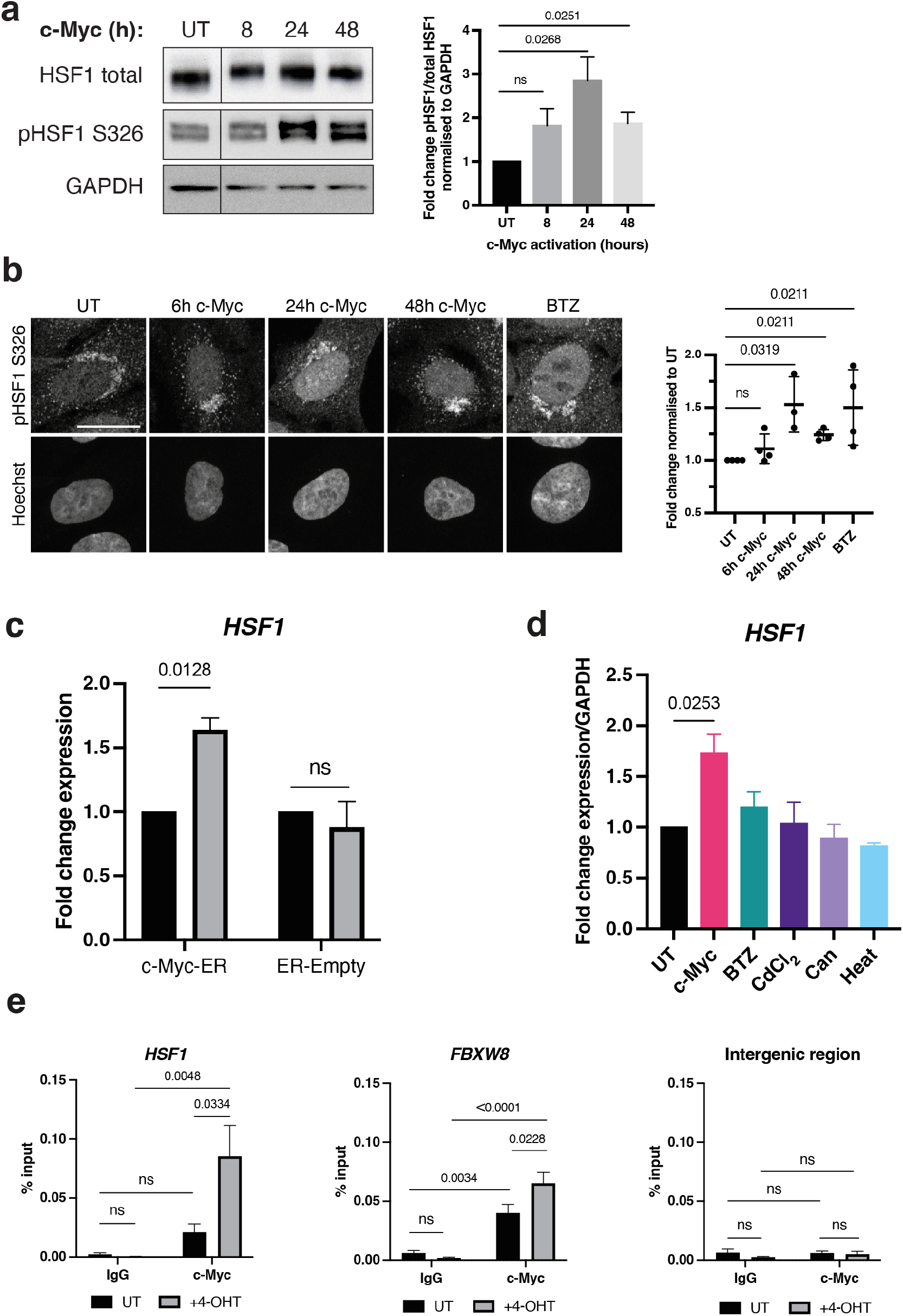
HSF1 is upregulated upon c-Myc activation post-translationally and transcriptionally. **(a)** c-Myc-ER cells were untreated (UT) or had c-Myc activated for the indicated timepoints before protein lysates were collected for Western blot, representative blot shown. Fold change values were calculated for n=3 independent experiments, error bars are SEM and unpaired Student’s t-test was used for statistical analysis. **(b)** c-Myc-ER cells were UT, had c-Myc activated for the indicated timepoints, or BTZ treatment for 6h before being fixed and stained for immunofluorescent staining of pHSF1 S326. Representative images shown, scale bar = 20*μ*m. Fold change values were calculated from n=3 (24h c-Myc) or 4 (all other samples) independent experiments, error bars are SEM and statistical analysis performed using Mann-Whitney test. **(c)** c-Myc-ER and ER-Empty cells were UT or had 24h 4-OHT treatment before samples were collected for RT-qPCR, n=3, error bars are SEM and 2-way ANOVA was used for statistical analysis. **(d)** c-Myc-ER cells were UT or had 24h of c-Myc activation, 6h BTZ, 6h cadmium chloride (CdCl_2_), 6h canavanine (Can), or 1h of heat shock before samples were collected for RT-qPCR. Error bars are SEM, n=3 (UT, c-Myc, BTZ, CdCl_2_) and n=2 (Can, Heat), statistical analysis performed using unpaired Student’s t-test. **(e)** c-Myc-ER cells were UT or treated for 16h with 4-OHT before cells were collected for chromatin immunoprecipitation (ChIP). Samples were pulled down with antibody for IgG or c-Myc before qPCRs were run targeting the promoter region of HSF1, FBXW8 or an intergenic region. Statistical analysis was run on percentage input values using two-way ANOVA, error bars are SEM, n=4 independent experiments.

Whilst proteotoxic stress induced HSF1 activation depends on post-translational modifications, increased gene expression of HSF1 has been observed in many cancers, suggesting that oncogenic signalling might be able to upregulate HSF1 directly. We wondered if activation of c-Myc induces transcriptional upregulation of HSF1. Using RT-qPCR we show that transcript levels of HSF1 are significantly increased upon 24h of 4-OHT treatment in c-Myc-ER cells, but not in ER-Empty cells (Fig. 2c). To establish if the transcriptional upregulation of HSF1 is c-Myc specific, rather than proteotoxic stress-induced, we compared the transcriptional response to c-Myc activation with that of several established proteotoxic stress-inducing treatments. Whilst, as expected, all tested treatments increase HSF1 S326 phosphorylation and induction of the HSF1-target HSPA1A (Supplementary Fig. S2a), we found that transcriptional upregulation of HSF1 was unique to c-Myc activation, with none of the traditional proteotoxic stress-inducers causing an increase in HSF1 mRNA levels (Fig. 2d). This suggests that the transcriptional upregulation of HSF1 is c-Myc-dependent.

The promoter region of HSF1 contains an E-box c-Myc binding site (Supplementary Fig. S2b) suggesting that transcriptional upregulation of HSF1 might be directly through c-Myc. To establish if HSF1 is a direct target of c-Myc, we assessed if c-Myc binds to the promoter region of HSF1 using chromatin immunoprecipitation (ChIP). We observe significant enrichment of the promoter region of the well-established c-Myc target FBXW8 (39), by anti-c-Myc antibody pulldown compared to IgG control, in both the uninduced (UT) and c-Myc induced (+4-OHT) samples. We also find enrichment of the HSF1 promoter region in the c-Myc samples, but interestingly this is only significant in cells with oncogenic levels of c-Myc activation (Fig. 2e). No enrichment of negative control intergenic region was found. These data suggest that c-Myc can bind the HSF1 promoter and indicates that oncogenic levels of c-Myc significantly increases binding, promoting HSF1 transcription.

Together, our data suggests that c-Myc increases HSF1 activation in two ways, by increasing post-translational modification of HSF1, likely due to c-Myc-induced proteotoxic stress, and by direct transcriptional upregulation, establishing a direct link for HSF1 activation by oncogenic signalling.

### HSF1 is required for a functioning DDR, through the HSP90AA1-dependent stabilisation of *γ*H2AX

Our data shows that c-Myc-induced proteotoxic stress contributes to the accumulation of DNA damage and that c-Myc, directly and indirectly, activates HSF1 to induce tolerance to proteotoxic stress. This suggests that HSF1 has an importance role in preventing c-Myc-induced DNA damage. To test this, we performed a time-course of c-Myc activation in c-Myc-ER cells transfected with non-targeting control siRNA (siControl) or HSF1-targeting siRNA (siHSF1). Protein levels of *γ*H2AX were used as a readout of DNA damage levels. As previously observed, activation of c-Myc leads to an increase in *γ*H2AX, indicating DNA damage accumulation, in siControl cells (Fig. 3a). Unexpectedly, the c-Myc-induced *γ*H2AX accumulation is lost upon knockdown of HSF1, indicating lack of DNA damage signalling. Loss of c-Myc-induced *γ*H2AX upon HSF1 knockdown was also observed via immunofluorescence staining (Fig. 3b) and with an alternative siRNA against HSF1 (siHSF1_b) via Western blot (Supplementary Fig. S3a).

**Fig. 3.**
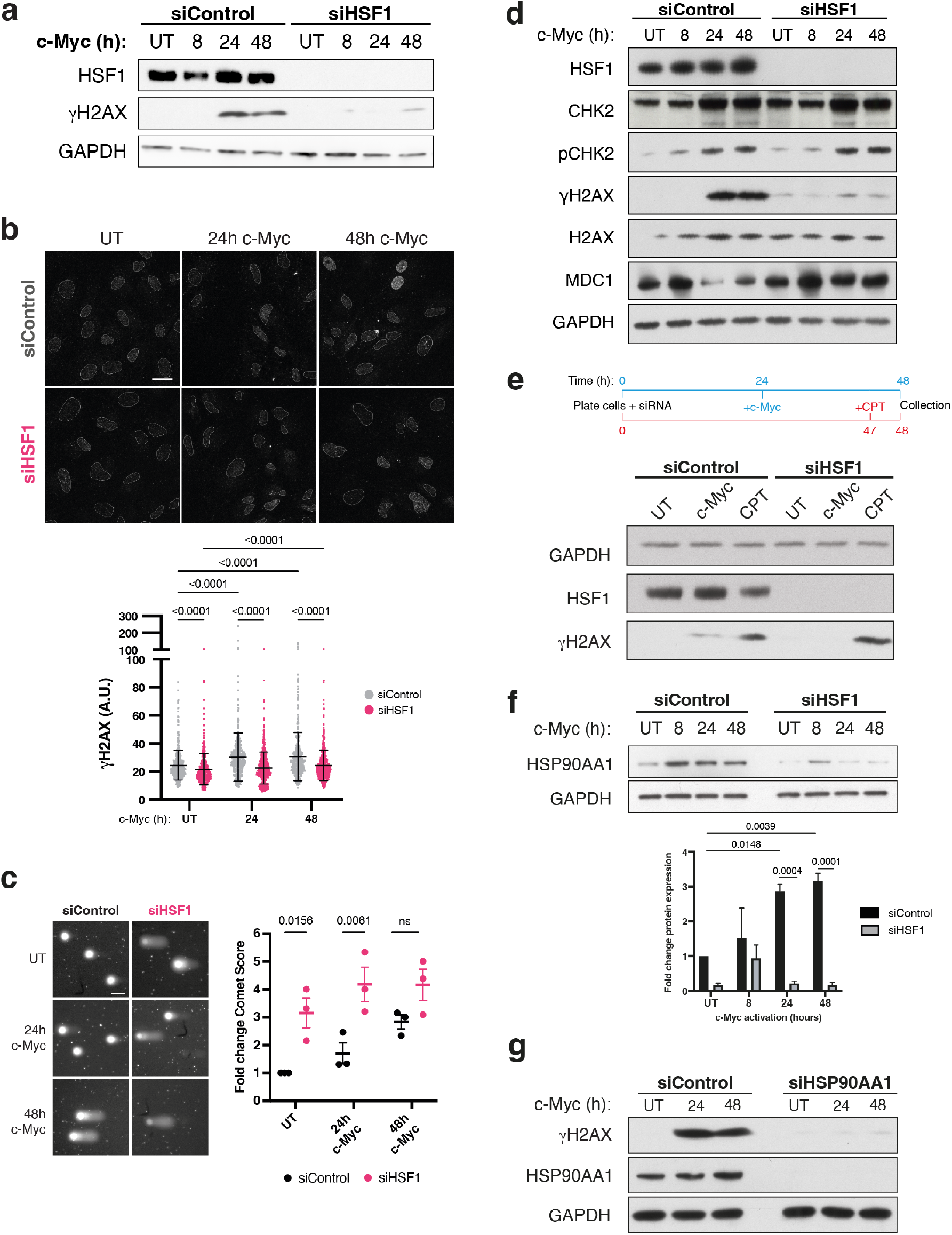
c-Myc-dependent upregulation of HSF1 is required for tolerance to proteotoxic stress-induced DNA damage and stabilisation of the DNA damage response component *γ*H2AX. **(a)** c-Myc-ER cells were transfected with siControl or siHSF1 siRNAs, with activation of c-Myc for the indicated timepoints before samples were collected for Western blot. Representative blot shown from n=3 independent experiments. **(b)** c-Myc-ER cells were transfected with siControl or siHSF1 siRNAs, with activation of c-Myc for the indicated timepoints before samples were collected for immunofluorescent staining of *γ*H2AX. Representative images shown, outlines indicate nuclear masks, scale bar = 20*μ*m. Intensity of *γ*H2AX was quantified for *≥*100 cells from each sample from n=3 independent experiments, error bars are SD and statistical analysis was performed using 2-way ANOVA. **(c)** c-Myc-ER cells were transfected with siControl or siHSF1 siRNAs and had c-Myc activated for the indicated timepoints before cells were collected for alkaline comet assay analysis. Scale bar = 50*μ*m. Fold change comet score for each condition shown, n=3, error bars are SEM. 2-way ANOVA was used for statistical analysis. **(d)** c-Myc-ER cells were transfected with siControl or siHSF1 siRNAs, with activation of c-Myc for the indicated timepoints before samples were collected for Western blot. Representative blot shown from n=3 independent experiments. **(e)** c-Myc-ER cells were transfected with siControl or siHSF1 siRNAs, with activation of c-Myc for 24h or CPT treatment for 1h before samples were collected for Western blot. Representative blot shown from n=3 independent experiments. **(f)** c-Myc-ER cells were transfected with siControl or siHSF1 siRNAs, with activation of c-Myc for the indicated timepoints before samples were collected for Western blot. Representative blot shown from n=3 independent experiments. Fold change expression was quantified from n=3 independent experiments for each sample, error bars are SEM and statistical testing performed using 2-way ANOVA. **(g)** c-Myc-ER cells were transfected with siControl or siHSP90AA1 siRNAs, with activation of c-Myc for the indicated timepoints before samples were collected for Western blot. Representative blot shown from n=3 independent experiments.

To test if this loss of *γ*H2AX was due to an absence of DNA damage or an impairment to DDR signalling we used the comet assay, which provides a direct readout of DNA damage without relying on upstream signalling markers of damage. In siControl c-Myc-ER cells, activation of c-Myc leads to an accumulation of DNA damage, as expected (Fig. 3c). Strikingly, knockdown of HSF1 in c-Myc-ER cells causes DNA damage already in untreated conditions and leads to a greater accumulation of DNA damage upon activation of c-Myc. This indicates that the loss of c-Myc-induced *γ*H2AX in cells treated with siHSF1 is due to a loss of DDR signalling.

The DDR depends on the initial activation of the checkpoint protein kinase ATM, which in addition to phosphorylating H2AX, phosphorylates and activates the effector kinase Chk2 and recruits the *γ*H2AX-binding partner MDC1 (40). The absence of c-Myc-induced histone H2AX phosphorylation upon knockdown of HSF1 could be due to loss or inactivity of any of these DDR-signalling proteins. We find that protein levels of the DDR effector kinase CHK2 and its activity by ATM-dependent phosphorylation at Thr68 (41, 42) is not affected by HSF1 knockdown in a time-course of c-Myc activation (Fig. 3d). Similarly, protein levels of the *γ*H2AX-binding partner MDC1 were not reduced upon knockdown of HSF1 and total H2AX levels were also unperturbed. These data suggest that the absence of *γ*H2AX upon siHSF1 treatment is not due to a loss of DDR protein levels or ATM-signalling.

An alternative reason for loss of *γ*H2AX could be that knock-down of HSF1 impairs the cells’ ability to stabilise *γ*H2AX. To investigate this, we tested if cells are able to phosphorylate H2AX upon acute induction of DNA damage by the topoisomerase I inhibitor camptothecin (CPT) in the absence of HSF1 (Fig. 3e). Acute DNA damage induction by CPT treatment, but not c-Myc activation, causes accumulation of *γ*H2AX in HSF1 knockdown cells. This indicates that HSF1 is not required for phosphorylation of H2AX, suggesting that an absence of c-Myc-induced *γ*H2AX in HSF1 knockdown cells is likely the result of a destabilisation event. An attractive candidate for the HSF1-dependent stabilisation of *γ*H2AX in the context of c-Myc activation is HSP90AA1. HSP90AA1 is a well-established target of HSF1 and colocalises with *γ*H2AX at sites of DNA damage (43). Our data show that c-Myc activation leads to an upregulation of HSP90AA1 protein and mRNA and that this is dependent on HSF1 expression, as c-Myc-induced HSP90AA1 levels are significantly decreased upon knockdown of HSF1 (Fig. 3f, Supplementary Fig. S3b). We next assessed if loss of HSF1-dependent accumulation of HSP90AA1 could be at the basis of the loss of c-Myc-induced *γ*H2AX accumulation, as seen in HSF1 knockdown cells. Knockdown of HSP90AA1 recapitulates the loss of c-Myc-induced *γ*H2AX, suggesting that the absence of c-Myc-induced *γ*H2AX in HSF1 knockdown cells is the likely consequence of a decrease in HSP90AA1 expression (Fig. 3g). Together, these data suggest that c-Myc-dependent upregulation of HSF1 is important for both preventing proteotoxic stress-induced DNA damage, as well as the HSF1/HSP90AA1-dependent maintenance of *γ*H2AX, required to initiate DDR signalling for repair of c-Myc-induced DNA damage.

### HSF1 prevents c-Myc-induced genomic instability to allow cell survival

Our data indicates that c-Myc-induced proteotoxic stress causes DNA damage, and that c-Myc-dependent upregulation of HSF1 is important to prevent and repair DNA damage. Next, we established the importance of HSF1 in preventing c-Myc induced genotoxic stress and genome instability. First, we established accumulation of 53BP1 foci, a central mediator and effector of the DDR coordinating DSB repair (44), upon activation of c-Myc in the presence and absence of HSF1. We find that while knockdown of HSF1 alone can cause an increase in 53BP1 foci, c-Myc-induced foci are substantially increased by HSF1 knockdown (Fig. 4a). This demonstrates the importance of c-Myc-induced upregulation of HSF1 for genotoxic stress tolerance. DNA damage left unresolved prior to entry into mitosis can lead to several defects, including the formation of micronuclei (45), which can be assessed by performing immunofluorescence imaging (Fig. 4b). Firstly, c-Myc activation increases micronuclei formation in both siControl and siHSF1 cells, as expected. Interestingly, we found that knockdown of HSF1 alone does not cause an increase in micronuclei formation. However, upon activation of c-Myc, the percentage of cells with micronuclei is significantly increased in siHSF1 cells compared to siControl, demonstrating a role for HSF1 for preventing c-Myc-induced genomic instability. Together these data suggest that whilst HSF1 is generally required to prevent genotoxic stress, only in cells experiencing oncogenic c-Myc activity is it necessary to prevent the accumulation of genomic instability.

**Fig. 4.**
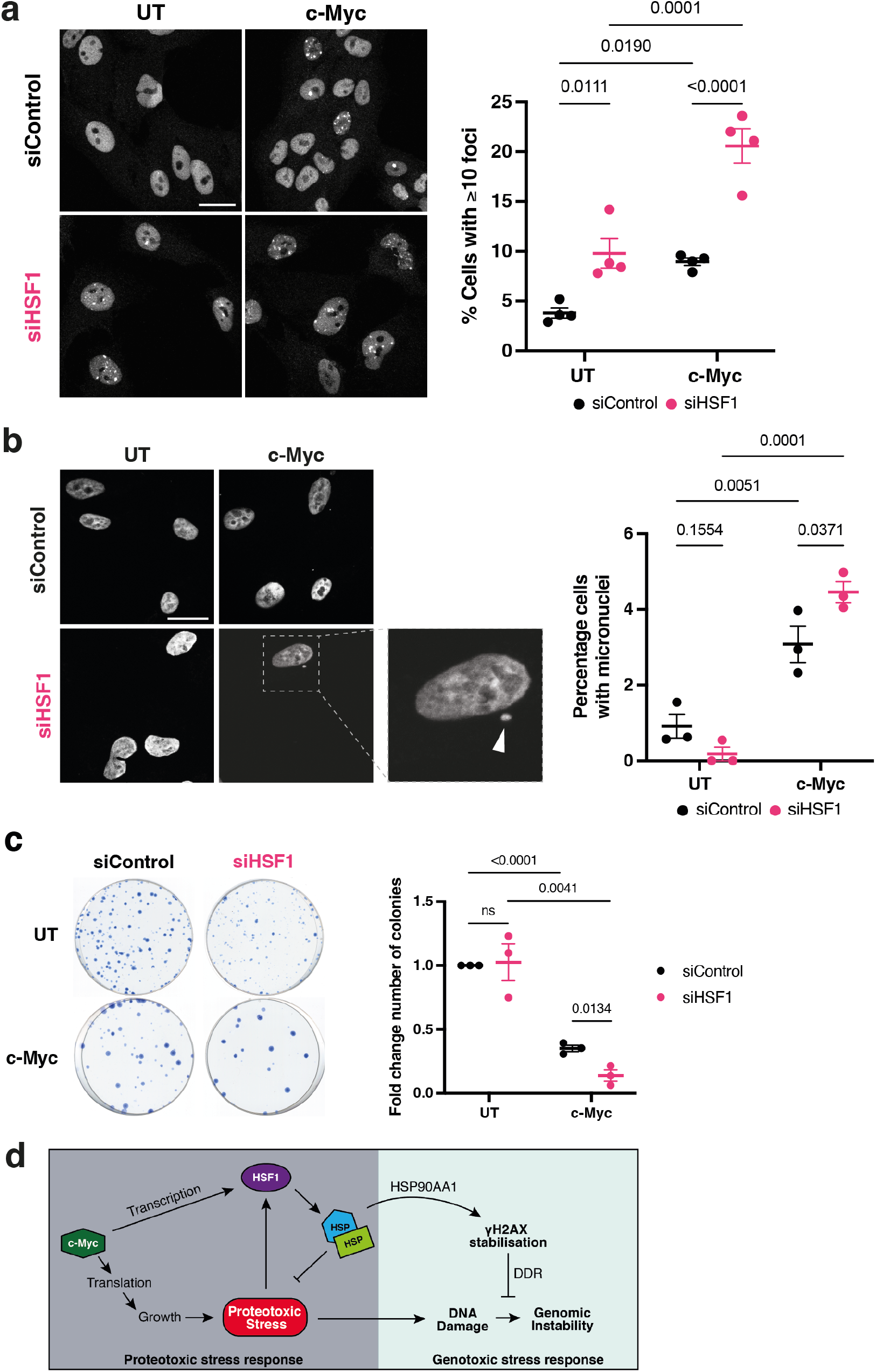
c-Myc increases tolerance to proteotoxic stress and DNA damage through HSF1. **(a)** c-Myc-ER cells were transfected with siControl or siHSF1 siRNAs and cells were either untreated or had c-Myc activated for 24h before cells were collected for immunofluorescent staining of 53BP1 foci. Representative images shown, scale bar = 20*μ*m. The percentage of cells with *≥*10 foci was quantified for n=4 independent experiments, error bars are SEM, statistical analysis performed using 2-way ANOVA. **(b)** c-Myc-ER cells were transfected with siControl or siHSF1 siRNAs and cells were either untreated or had c-Myc activated for 24h before cells were collected for immunofluorescent staining of DNA with Hoechst. Representative images shown, scale bar = 20*μ*m. The percentage of cells with micronuclei was quantified for n=3 independent experiments, error bars are SEM, statistical analysis performed using 2-way ANOVA. **(c)** c-Myc-ER cells were transfected with siControl or siHSF1 siRNAs and were either untreated or had c-Myc activated for 24h before cells were replated at low density and left to grow for 7 days. Colonies were stained with methylene blue and counted to calculate the fold change in colony number from n=3 representative experiments. Error bars are SEM and statistical analysis performed using 2-way ANOVA. **(d)** Proposed model for how c-Myc increases tolerance to proteotoxic stress and DNA damage through HSF1.

Since HSF1 seems to be required to prevent c-Myc-induced genomic instability, we next tested if HSF1 is important for cell viability in cells experiencing oncogenic c-Myc activity using colony formation assays. In line with our micronuclei data, knockdown of HSF1 in untreated c-Myc-ER cells does not lead to a significant reduction in colony number (Fig. 4b, Supplementary Fig. S4a). However, in cells experiencing oncogenic c-Myc activity colony numbers are significantly decreased upon loss of HSF1, demonstrating the importance of HSF1 for the survival of c-Myc cells. 4-OHT addition to ER-Empty cells does not cause a reduction in colony number, demonstrating that the loss of cell viability observed in c-Myc-ER cells is c-Myc-dependent (Supplementary Fig. S4b).

Together, these data suggest a central role for HSF1 in the tolerance to c-Myc-induced proteotoxic and genotoxic stress, whereby loss of HSF1 increases genomic instability and reduces cell viability in the context of c-Myc activation.

## Discussion

Together, our data supports a model whereby oncogenic c-Myc activity increases tolerance to proteotoxic and genotoxic stress to prevent genomic instability through regulation of HSF1 (Fig. 4d). We show that increased c-Myc activity upregulates HSF1, through direct transcriptional regulation, and as a consequence of c-Myc-induced proteotoxic stress. This is not only important to prevent proteotoxic stress-induced DNA damage but is also required for DDR signalling through *γ*H2AX. Whilst HSF1’s role in the tolerance to oncogene-induced proteotoxic stress has been well documented, our study uncovers an important role for c-Myc-induced upregulation of HSF1 to prevent genomic instability, ensuring that c-Myc-induced growth and proliferation is compatible with cell survival.

Our data indicates that DNA damage is in part driven through an increase in c-Myc-dependent proteotoxic stress, revealing a potentially important role for oncogene-induced proteotoxic stress in the generation of genomic instability. Our work establishes a key role for HSF1 in not only preventing proteotoxic stress-induced DNA damage, but also tolerance to DNA damage through DDR signalling via *γ*H2AX, linking the proteotoxic and genotoxic stress responses. Oncogenic signalling has previously been shown to activate HSF1 (14–19, 46). Our work demonstrates for the first time the direct involvement of oncogenic c-Myc in HSF1 upregulation (Fig. 4d). With c-Myc downstream of most oncogenic signalling that drives growth and proliferation, it establishes a mechanism by which c-Myc-dependent regulation of HSF1 can facilitate the increased reliance of cancer cells on the proteotoxic stress response. This provides a mechanism by which oncogenic signalling can upregulate HSF1 directly to adapt to proteotoxic and genotoxic stress resulting from sustained growth and proliferation.

Overall, our work suggests that c-Myc-induced proteotoxic stress can contribute to the accumulation of DNA damage and that the cellular transcriptional response, via HSF1, is required to prevent this. HSF1 promotes transcription of heat shock proteins that guide protein folding, trafficking and disaggregation. This is required to lower general proteotoxic stress levels but is also important for the correct function of proteins involved in preventing or repairing DNA damage (47, 48). More generally, our work suggests that proteotoxic stress can contribute to the rapid genome evolution observed in tumours, generating intra-tumour heterogeneity, which is implicated in cancer multidrug resistance and adverse clinical outcomes (49).

## Materials and Methods

### Cell Culture

RPE1-TetON c-Myc-ER cells were generated as described previously (30), and will be hereafter described as RPE1 c-Myc-ER cells. Briefly, RPE1 hTERT cells were retrovirally infected with pBABE c-Myc-ER plasmid (Addgene, 19128), infected cells were selected in 5*μ*g/ml puromycin and the surviving polyclonal population was used in further assays. RPE1-TetON ER-Empty cells (hereafter RPE1 ER-Empty) were generated as above, using a control pBABE-ER plasmid. Cells were cultured at 37°C, 5% CO_2_in DMEM/F12 phenol red-free media (Gibco, 21041) supplemented with 7% charcoal treated Fetal Bovine Serum (FBS) (Sigma, F7524), 1% Penicillin-Streptomycin (P/S) (Gibco, 15140), 3.5% sodium bicarbonate (Gibco, 25080) and initially selected in 2*μ*g/ml Puromycin when thawing a new vial of cells. Charcoal treated FBS was prepared by addition of 0.5g Charcoal (Sigma, C9157) and 0.05g Dextran (Sigma, D8906) to 500ml FBS, stirring for 2.5h at 37°C, then filtering through 0.2*μ*M vacuum filters (Merck Millipore, SCGVU05RE).

### Cell Treatments

To activate c-Myc, Hydroxytamoxifen (4-OHT) (Sigma, H7904) was used at a final concentration of 100nM in complete media for the indicated timepoints. Bortezomib (Stratech Scientific, S1013-SEL) was used at a final concentration of 20nM in 0.1% DMSO for 6 or 24 hours, as indicated, before collection. Cadmium chloride (CdCl_2_) (Sigma, 202908) was used at a final concentration of 10*μ*M in H_2_O for 6 hours before collection. Canavanine (Sigma, C9758) was used at a final concentration of 20mM in H_2_O for 6 hours before collection. Torin1 (Merck Millipore, 475991) was used at a final concentration of 25nM in 0.1% DMSO for 24 hours before collection. Heat shock was performed by sealing tissue culture dishes with parafilm and submerging them in a 42°C water-bath for 1 hour. After the heat-shock, dishes were wiped with 70% ethanol and dried with tissue paper before being placed back in a 37°C, 5% CO_2_ incubator for 5 hours of recovery. Camptothecin (Sigma, C9911) was used at a final concentration of 2*μ*M in 0.1% DMSO for 1 hour before collection. Hydroxyurea (Sigma, H7904) was used at a final concentration of 0.1mM in H_2_O for 4 hours before collection.

### Transfection

For siRNA transfection, Lipofectamine RNAiMAX (Thermo Fisher, 13778075) was used following the manufacturer’s reverse transfection protocol, with all dilutions made in OptiMEM no phenol red (Gibco, 11058021). A final concentration of 20nM siRNA was used in all transfections. Cells were plated on top of transfection complexes at a concentration of 4×10^4^ cells/ml in P/S-free media and cultured overnight. The next morning, culture media was replaced with complete media and cultured for the indicated timepoints. Non-targeting control siRNA (Dharmacon, Horizon Discovery, D-001810-01-20) was used as a negative control in all experiments. Custom siRNA targeting HSF1 was synthesised by IDT with a Sense sequence of 5’-CCUGAAGAGUGAAGACAUAUU-3’ and anti-Sense sequence of 3’-GGACUUCUCACUUCUGUAUAA-5’.The following siRNAs were pre-designed by Dharmacon, Horizon Discovery: siHSF1_b (L-012109-01-0005), ATF4 (L-005125-00-0005), HSP90AA1 (L-005186-00-0005).

### Western Blot

Cell extracts were prepared in RIPA buffer (Tris-HCL pH 7.5 20mM, NaCl 150mM, EDTA 1mM, EGTA1mM, NP40 1%, NaDOC 1%) containing protease inhibitor cocktail (Sigma, P8340) and phosphatase inhibitor cocktails 2 (Sigma P5726) and 3 (Sigma P0044). Protein concentration was determined using the Bradford assay (BioRad, 500-0006) following the manufacturer’s protocol. Equal amounts of protein were loaded into Nu-Page Novex 4-12% Bis-Tris protein gels (Invitrogen, NP0322) and electrophoresis run in MOPS buffer (Invitrogen, NP0001). Protein was wet transferred to nitrocellulose membranes (Sigma, GE10600001) in transfer buffer (25mM Tris base, 250mM glycine, 20% ethanol). Membranes were blocked for 1 hour in either 5% milk or 5% Bovine Serum Albumin (BSA) (Sigma, A7906) diluted in PBS/0.2% Tween, following antibody-supplier instructions. Membranes were then incubated overnight at 4°C in primary antibodies diluted in 5% milk or 5% BSA PBS/0.2% Tween. Membranes were washed in PBS/0.2% Tween and incubated in secondary HRP antibody for 1 hour at room temperature in 5% milk or 5% BSA PBS/0.2% Tween. HRP was visualised using Luminata Crescendo HRP substrate (Merck, WBLUR0100), developed onto Amersham Hyperfilm (GE Healthcare Life Sci-ences, 28906836) with a XOGRAF Compact X4 film processor. Antibodies used were as follows: ATF4 (Cell Signaling Technologies, 11815), CHK2 (Cell Signaling Technologies, 2662), CHK2 phospho-S345 (Cell Signaling Technologies, 2661), Cyclin E (Invitrogen, MA5-32358), GAPDH (GeneTex, GTX627408), H2AX (Cell Signaling Technologies, 2595), H2AX phospho-S139 (Cell Signaling Technologies, 9718), HSF1 (Abcam, ab52757), HSF1 phospho-S326 (Abcam, ab76076), HSP90AA1 (Abcam, ab79848), HSPA1A (Abcam, ab5439), MDC1 (Abcam, ab241048),Vinculin (Abcam, ab129002), anti-mouse-HRP (Fisher Scientific, PA1-74421), anti-rabbit-HRP (Fisher Scientific, PA1-31460).

### RT-Qpcr

RNA was extracted using a Qiagen RNeasy Plus Mini Kit (Qiagen, 74134) following the manufacturer’s protocol. The reaction was carried out using One Step Mesa Blue mastermix (Eurogentec, 032XNR), supplemented with Euroscript Reverse Transcriptase/RNase inhibitor (Eurogentec, RT-0125-ER) for reverse transcription of RNA. Reactions were carried out in a total volume of 14*μ*l per well, with 7*μ*l of Mesa Blue (supplemented with 0.035*μ*l Euroscript reverse transcriptase), 1.5*μ*l of 5*μ*M forward primer, 1.5*μ*l 5*μ*M reverse primer and 4*μ*l of 20ng/*μ*l RNA product. Each sample was run in triplicate.The qPCR reaction was run on a BioRad CFX Connect machine. Reverse transcription qPCR steps were as follows: 30m at 48°C, 5m at 95°C, 40 cycles of 3s at 95°C and 45s at 60°C and melt curve analysis from 65-85°C with 10s at each temperature in increments of 1°C. Data was analysed using the △△Ct method to normalise transcript levels to the housekeeping gene GAPDH. Statistical analysis was performed on the ΔCt values. Primer sequences were as follows (5’-3’): *CDC45* FGAAGAGGAGATAGTGGAGCAAAC, R GCTGACGAT-GTCCCATGATATT; *eIF4E* F GGGAGGGTTGATTGCC-TAAA, R GACAAGTTGCCTGTGTGTTTATT; *GAPDH* F GAAATCCCATCACCATCTTCCAGG, R GAGCC-CCAGCCTTCTCCATG; *HSF1* F GGAGTCCATAGCATC-CAAGTG, R CCTCCACCCCTGAAAAGTG; *HSP90AA1* F TGAACTGGCGGAAGATAAAGA, R GCCTTCCT-GACCAAGCTGT; *MDM2* F GATCCAGGCAAATCT-GCAATAC, R TGGTCTAACCAGGGTCTCTT; *PAICS* F GACAAGAGGAGAAGCAGAGATTAG, R TGAG-GTATTAGTCGCATTT.

### Chromatin Immunoprecipitation PCR (ChIP-PCR)

Cells were cultured in 15cm dishes for 16h with or without addition of 4-OHT. Cells were then washed with PBS before crosslinking was performed via addition of 1% formaldehyde for 10m at room temperature. The crosslinking was quenched by addition of 1.25M glycine for 10m at room temperature. Cells were then collected in PBS and spun down, with the cell pellets resuspended in 1ml ice cold buffer A (100mM HEPES pH 8, 100mM EDTA, 5mM EGTA, 2.5% Triton-X100) and rocked at 4°C for 10m. Another wash was then repeated using ice cold buffer B (100mM HEPES pH 8, 2M NaCl, 100mM EDTA, 5mM EGTA, 0.1% Triton-X 100) before cells were resuspended in ice cold ChIP buffer (25mM Tris-HCl pH 8, 2mM EDTA, 15 mM NaCl, 1% Triton-X 100, 0.1% SDS). Cells were sonicated with a Bioruptor Pico machine for 30s on, 30s off for 20m at maximum output. Following sonication, the lysates were spun at maximum speed for 15m at 4°C. Protein A solution was made my resuspending beads in ChIP buffer with 1*μ*g/*μ*l BSA and 0.25*μ*g/*μ*l ssDNA to create a 50% slurry, and rocking for 10m at 4°C. The supernatant was transferred into fresh eppendorf tubes and pre-cleared by adding the blocked protein A solution and rocking at 4°C for 2h. Cleared soluble chromatin was centrifuged for 4m, 4000rpm, 4°C and the supernatant transferred into fresh eppendorf tubes. At this point, 1% of the solution was reserved to use as an input. The remaining soluble chromatin was incubated overnight at 4°C with 3*μ*g of either c-Myc antibody (anti-c-Myc, Abcam, ab56) or mouse IgG antibody (normal mouse IgG, Santa Cruz Biotechnology, sc-2025). The next morning, 20*μ*l of freshly made protein A beads were added to the chromatin and rocked for a further 2h at 4°C. The beads were then spun down for 2m at 2000rpm and washed in ChIP buffer, wash buffer 1 (25mM Tris-HCl pH 8, 2mM EDTA, 500mM NaCl, 1% Triton-X 100, 0.1%SDS), wash buffer 2 (250mM LiCl, 1% NP40, 1% NaDOC, 1mM EDTA, 10mM Tris-HCl pH 8), and twice in TE pH 8. The samples were then spun and the pellets resuspended in elution buffer (1% SDS, 100mM NaHCO_3_). The samples were then incubated overnight at 65°C in order to reverse crosslinking. The following morning, RNAse A was added to each sample for a further 30m at 65°C, before samples were purified using QIAquick PCR Purification kit (Qiagen) and diluted in ddH2O. The qPCR reaction was carried out as above with 4*μ*l of DNA product. Each sample was run in triplicate. The qPCR reaction was run on a BioRad CFX Connect machine. ChIP-qPCR steps were as follows: 5m at 95°C, 40 cycles of 3s at 95°C and 45s at 60°C and melt curve analysis from 60-85°C with 10s at each temperature in increments of 1°C. Data was analysed using the percent input method. Primer sequences were as follows (5’-3’): *FBXW8* F GTGATAGGCAGCAGAGCTGA, R TGTACGCACGTG-GTGGTC; *HSF1* F CAGTCCGCTCTGCCTGAGAC, R CCCGCGCCCAGTGAATC; Intergenic F ATGTCAGGCC-CATGAACGAT, R GCATTCATGGAGTCCAGGCTTT.

### Immunofluorescence

Cells were plated on coverslips precoated with fibronectin (Sigma, F1141). Cells were extracted to remove protein not bound to chromatin. For extraction, cells were incubated in ice cold PBS 0.2% Triton X-100 (Sigma, T8787) for 1m, then fixed with 4% PFA for 20m. If extraction was not required, cells were fixed in 4% PFA for 20m, rinsed with PBS 0.2% Tween and then permeabilised in PBS 0.2% Triton X-100 for 5m. Coverslips were then blocked in 1% BSA PBS 0.2% Tween for 1h at room temperature and incubated in primary antibody in blocking solution, overnight at 4°C. Coverslips were then washed with PBS 0.2% Tween and incubated in secondary antibody for 1h at room temperature. Following PBS 0.2% Tween washes, coverslips were incubated in Hoechst 3342 trihydrochloride, trihydrate (Invitrogen, H3570, 1:10,000) for 5m at room temperature and mounted on slides with Fluoroshield (Sigma,F6182). Coverslips were left to dry and then sealed with clear nail varnish. Images were acquired using a Leica TCS SPE2 confocal microscope with a 63x (1.3 NA) or 40x (1.15 NA) oil objective lens and processed in Fiji. Antibodies used were 53BP1 (Santa Cruz, sc-22760), H2AX phospho-S139 (Cell Signaling Technologies, 9718), HSF1 phospho-S326 (Abcam, ab76076), Alexa Fluor 488 anti-mouse IgG (Life Technologies, A11029), Alexa Flour 647 anti-rabbit IgG (Life Technologies, A21244).

### Proteostat Protein Aggregation Assay

For analysis of protein aggregation, the Proteostat Aggresome detection kit (Enzo Life Sciences, ENZ-51035-0025) was used. Cells were trypsinised and the pellet washed in PBS. Cells were fixed in 4% PFA for 30m at room temperature, pellet washed in PBS and then permeabilised in PBS/0.05% Triton X-100 for 30m on ice. The pellet was then washed in PBS and resuspended in freshly made Proteostat Aggresome Detection Reagent and incubated for 30m at room temperature. Samples were measured on a BD LSRII flow cytometer in the FL3 channel using DIVA software and analysed with FlowJo. Cells were gated firstly according to size using a forward scatter (FSC) vs side scatter (SSC) plot. Cells were secondly gated to exclude doublets and cell clumps using a forward scatter area (FSC-A) vs height (FSC-H) plot.

### Comet assay

The Trevigen CometAssay Reagent Kit for Single Cell Gel Electrophoresis Assay (Trevigen, 4250-050-K) was used. Cells were collected by trypsinisation and resuspended in ice-cold Ca_2_^+^ and Mg_2_^+^ free PBS (Sigma, D8537) and processed following the manufacturer’s protocol for Alkaline Comet Assay. Lysis was performed at 4°C overnight and electrophoresis was run at 1V/cm and a current of 300mA for 40m at 4°C. DNA was stained with 10*μ*g/ml of propidium iodide. Images of comets were captured on a Zeiss Axio Imager widefield microscope with 10x (0.3 NA) air objective lens. Scoring of comets was performed manually using established grading criteria as outlined previously (50).

### Colony Formation Assay

After treatment, cells were harvested by trypsinisation and counted with a haemocytometer. 150 cells per treatment were re-plated per well of a 6-well plate and left to incubate at 37°C, 5% CO_2_ for 7 days. Cells were fixed in 70% ethanol and colonies stained with 1% Methylene Blue (Sigma, M4159) for 20m at room temperature. Excess stain was washed with ddH_2_O and stained colonies were counted manually.

### Statistics

Graphpad Prism software was used for statistical tests. For Figures 1a, 1c, 1e, 2a, 2d, S1b, S1c, S1d, S2a, and S4b, significance was determined using the unpaired two-tailed Student’s t-test. For Figure 2b significance was determined using the unpaired two-tailed Mann-Whitney test. For Figures 2c, 2e, 3b, 3c, 3f, 4a, 4b, 4c, S1a, S1f, S3b, and S4a significance was determined using a two-way ANOVA with Sidak’s multiple comparison test.

## ACKNOWLEDGEMENTS

We are grateful to Dr. J. Labbadia for advice. This work was supported by core funding to the MRC-UCL University Unit (MC_U12266B) and funded by R.d.B.’s Cancer Research UK Programme Foundation Award (CRUK CDF: C63833/A25729).

## Supplementary Information

**Fig. S1.**
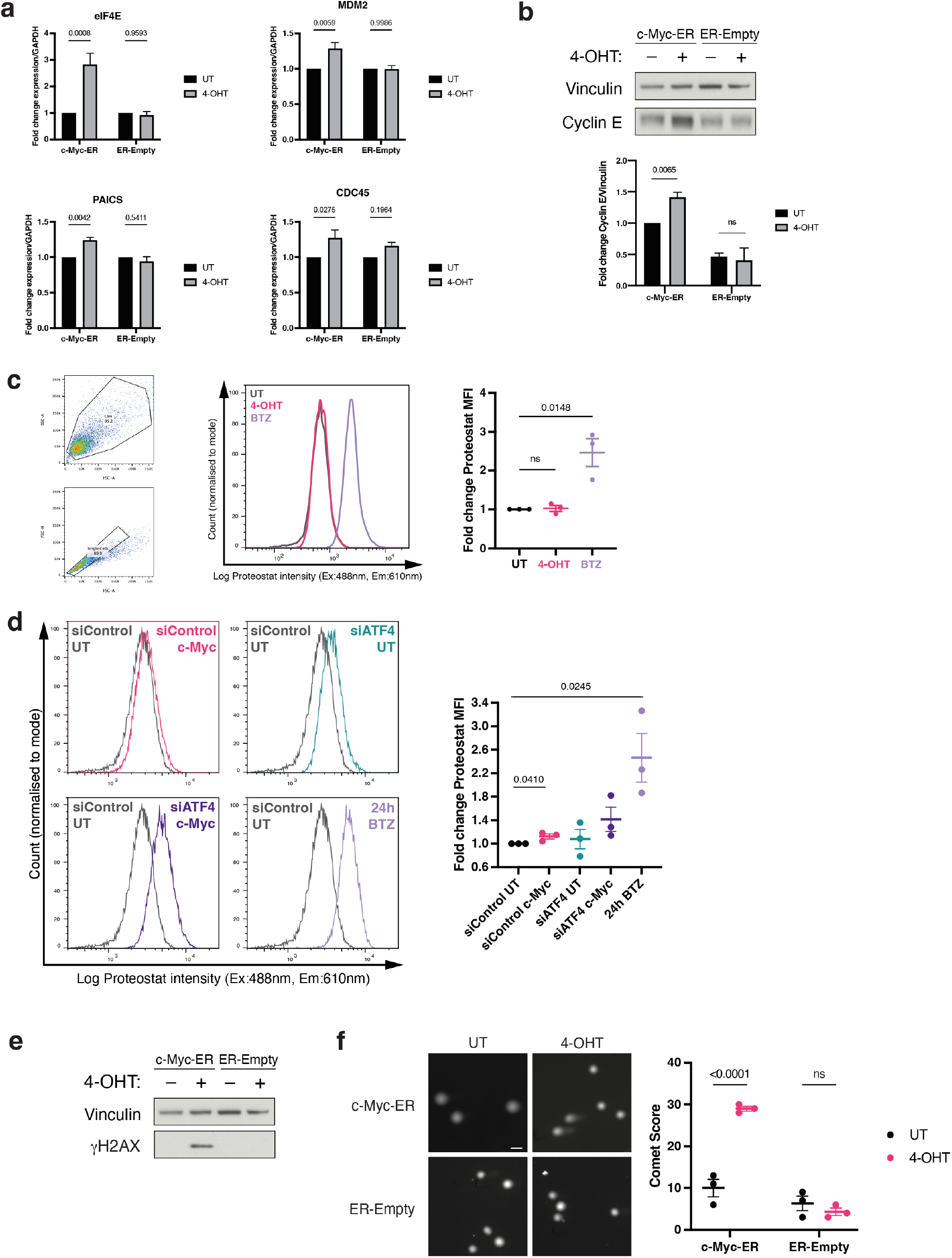
c-Myc-induced protetotoxic stress generates DNA damage. **(a)** c-Myc-ER or ER-Empty cells were untreated or had 4-OHT treatment for 24h, before cells were collected for RT-qPCR. Error bars are SEM, n=3, statistical testing performed using 2-way ANOVA. **(b)** c-Myc-ER or ER-Empty cells were untreated or had 24h of 4-OHT treatment before protein lysates were collected for Western blot, representative blot shown. Error bars are SEM, n=3, statistical testing performed using unpaired Student’s t-test. **(c)** ER-Empty cells were untreated or had 24h 4-OHT treatment before cells were fixed and stained with Proteostat for flow cytometry analysis, gating strategy and representative histograms shown. Fold change Proteostat median fluorescent intensity (MFI) was calculated for each sample, n=3, error bars are SEM. Unpaired Student’s t-test was used for statistical analysis. **(d)** c-Myc-ER cells were transfected with either siControl or siATF4 siRNA and had c-Myc activated for 24h or BTZ treatment for 24h, before cells were fixed and stained with Proteostat for flow cytometry analysis, representative histograms shown. Fold change proteostat median fluorescent intensity (MFI) was calculated for each sample, n=3, error bars are SEM. Unpaired Student’s t-test was used for statistical analysis.**(e)** c-Myc-ER or ER-Empty cells were untreated or had 24h 4-OHT treatment before protein lysates were collected for Western blot, representative blot shown, n=3.**(f)** c-Myc-ER or ER-Empty cells were untreated or had 24h 4-OHT treatment before cells were collected for alkaline comet assay analysis. Scale bar = 50*μ*m. Comet score for each condition shown, n=3, error bars are SEM. 2-way ANOVA was used for statistical analysis.

**Fig. S2.**
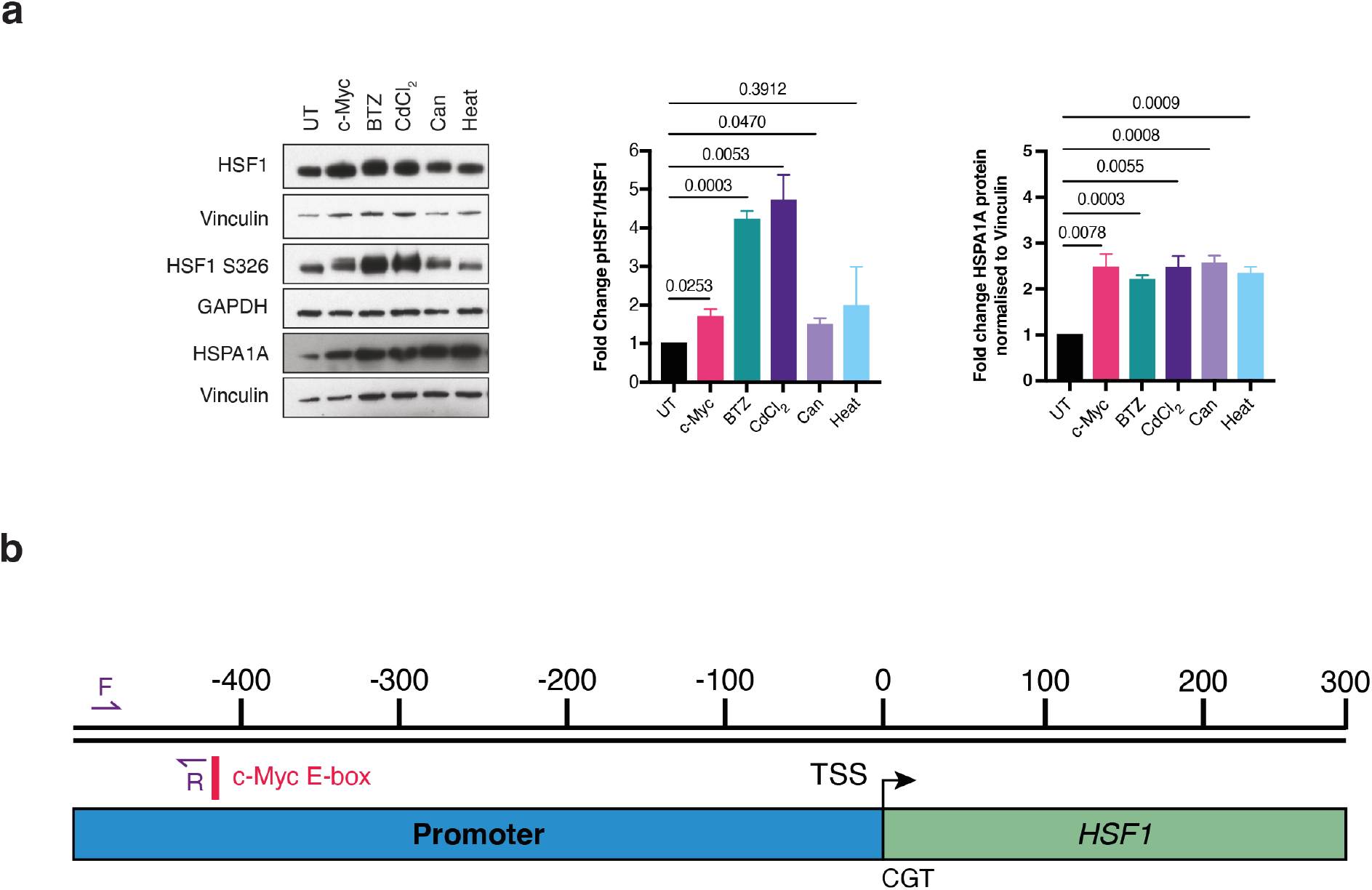
HSF1 is upregulated upon c-Myc activation post-translationally and transcriptionally. **(a)** c-Myc-ER cells were UT or had 24h of c-Myc activation, 6h BTZ, 6h cadmium chloride (CdCl2), 6h canavanine (Can), or 1h of heat shock before samples were collected for Western blot, representative blot shown. Fold change values calculated from n=3 independent experiments, error bars are SEM and statistical analysis performed using unpaired Student’s t-test. **(b)** Schematic for HSF1 ChIP-qPCR, showing forward (F) and reverse (R) primer binding regions, c-Myc E-box binding site and HSF1 transcription start site (TSS). Numbers indicate base pairs upstream (-) or downstream (+) of the TSS.

**Fig. S3.**
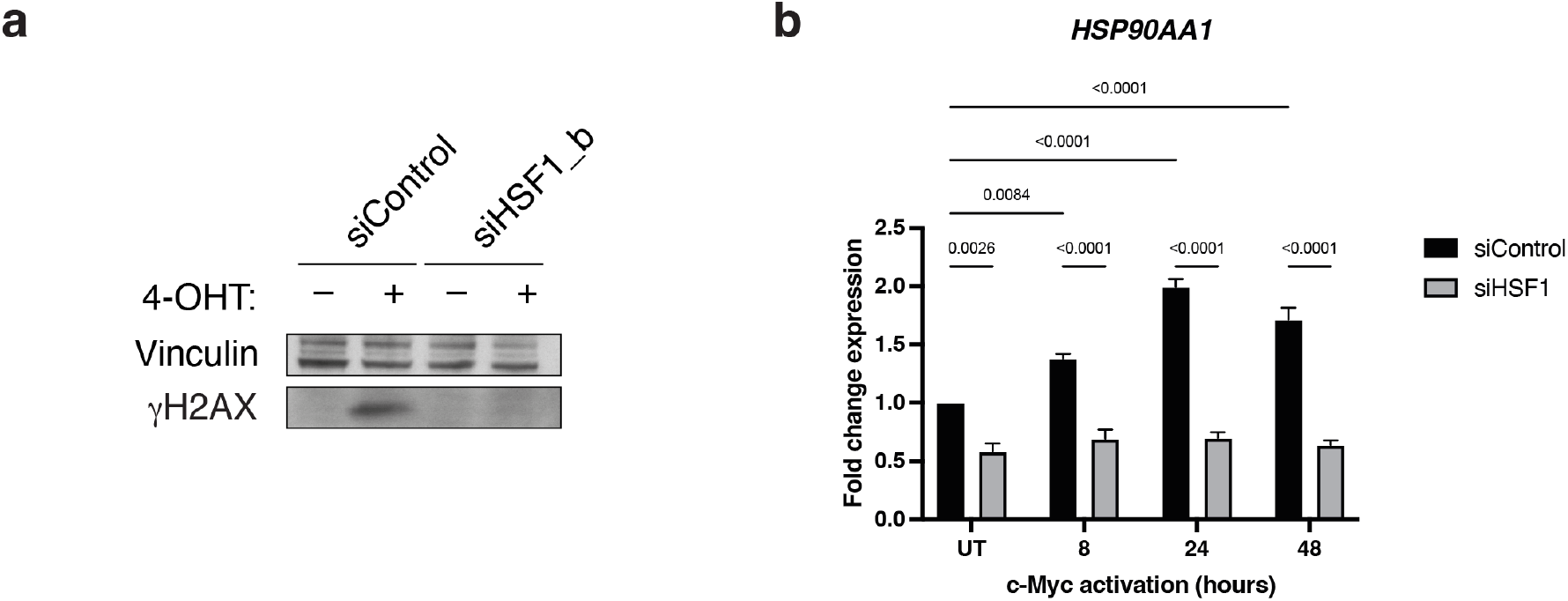
c-Myc-dependent upregulation of HSF1 is required for tolerance to proteotoxic stress-induced DNA damage and stabilisation of the DNA damage response component *γ*H2AX. **(a)** c-Myc-ER cells were transfected with siControl or siHSF1_b siRNAs with or without 24h of 4-OHT treatment before samples were collected for Western blot. Representative blot shown from n=2 independent experiments. **(b)** c-Myc-ER cells were transfected with siControl or siHSF1 siRNAs before c-Myc was activated for the indicated timepoints and samples collected for RT-qPCR. Error bars are SEM, n=3, statistical testing performed using 2-way ANOVA.

**Fig. S4.**
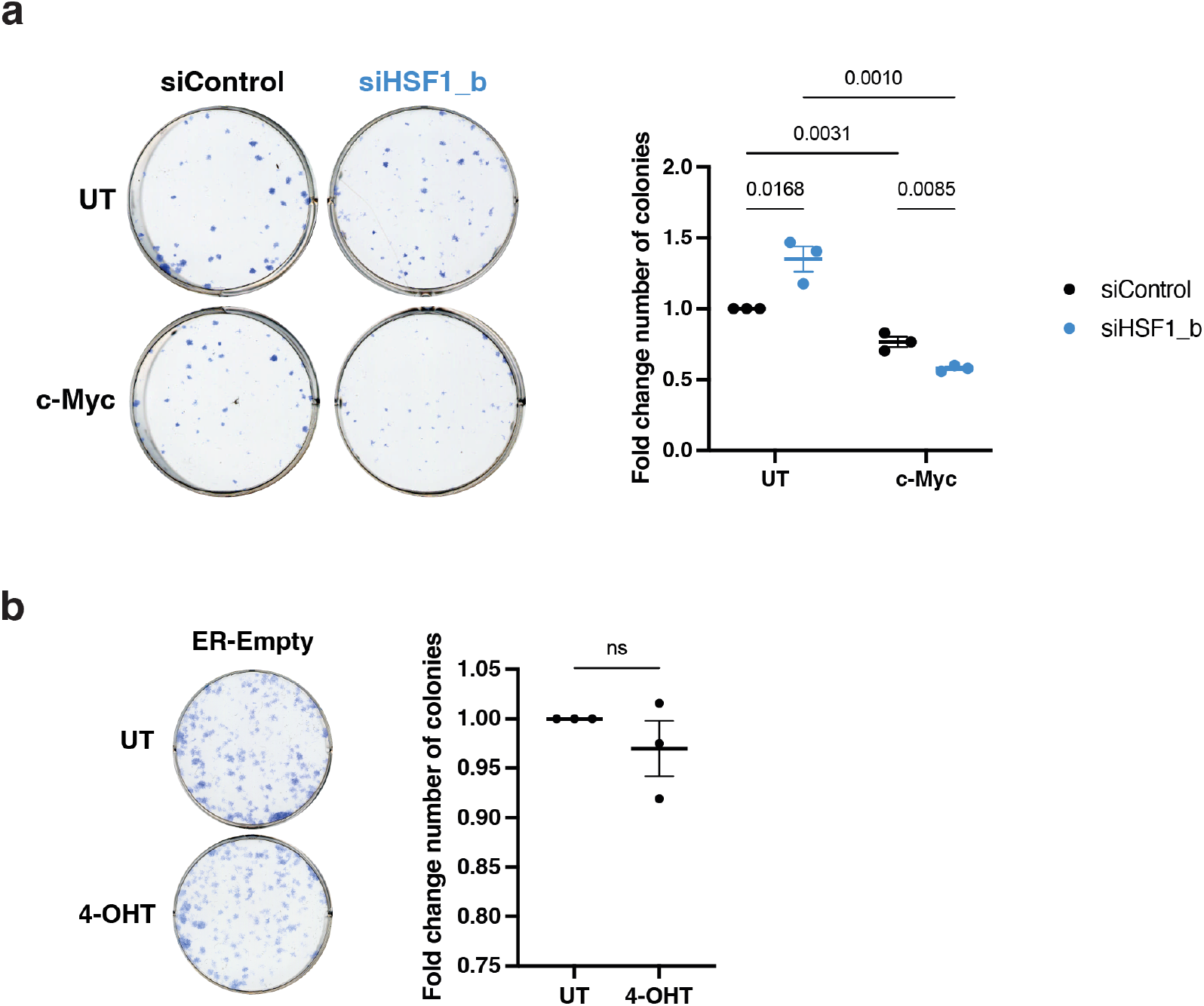
c-Myc increases tolerance to proteotoxic stress and DNA damage through HSF1. **(a)** c-Myc-ER cells were transfected with siControl or siHSF1_b siRNAs and were either untreated or had c-Myc activated for 24h before cells were replated at low density and left to grow for 7 days. Colonies were stained with methylene blue and counted to calculate the fold change in colony number from n=3 representative experiments. Error bars are SEM and statistical analysis performed using 2-way ANOVA. **(b)** ER-Empty cells were either untreated or had 24h 4-OHT treatment before cells were replated at low density and left to grow for 7 days. Colonies were stained with methylene blue and counted to calculate the fold change in colony number from n=3 representative experiments. Error bars are SEM and statistical analysis performed using unpaired Student’s t-test.

